# Sexual dimorphism of the synovial transcriptome underpins greater PTOA disease severity in male mice following joint injury

**DOI:** 10.1101/2022.11.30.517736

**Authors:** Rachel F. Bergman, Lindsey Lammlin, Lucas Junginger, Easton Farrell, Sam Goldman, Rose Darcy, Cody Rasner, Alia M. Obeidat, Anne-Marie Malfait, Rachel E. Miller, Tristan Maerz

## Abstract

**Objective:** To elucidate sex differences in synovitis, mechanical sensitization, structural damage, bone remodeling, and the synovial transcriptome in the anterior cruciate ligament rupture (ACLR) mouse model of post-traumatic osteoarthritis (PTOA).

**Methods:** Male and female 12-week-old C57Bl/6 mice were randomized to noninvasive ACLR or sham (n=9/sex/group/timepoint). Knee hyperalgesia, mechanical allodynia, and intra-articular MMP activity (via intravital imaging) were measured longitudinally. Trabecular and subchondral bone remodeling and osteophyte formation were assessed by μCT. Histological scoring of PTOA and synovitis and anti-MMP13 immunostaining was performed. Na_V_1.8-Cre;tdTomato mice were used to document localization and sprouting of nociceptors. Bulk RNAseq of synovium in sham, 7d, and 28d post-ACLR, and contralateral joints (n=6) assessed injury-induced and sex-dependent synovial gene expression.

**Results:** Male mice exhibited worse joint damage at 7d and 28d and worse synovitis at 28d, accompanied by greater MMP activity, knee hyperalgesia, and mechanical allodynia. Females had catabolic responses in trabecular and subchondral bone after injury, whereas males exhibited greater osteophyte formation and sclerotic remodeling of trabecular and subchondral bone. Na_V_1.8+ nociceptor sprouting in subchondral bone and medial synovium was induced by injury and comparable between sexes. RNAseq of synovium demonstrated that both sexes had similar injury-induced gene expression at 7d, but only female mice exhibited synovial inflammatory resolution by 28d, whereas males had persistent pro-inflammatory, pro-fibrotic, pro-neurogenic, and pro-angiogenic gene expression.

**Conclusion:** Worse overall joint pathology and pain behavior in male mice was associated with persistent activation of synovial inflammatory, fibrotic, and neuroangiogenic processes, implicating persistent synovitis in driving sex differences in murine PTOA.

## Introduction

The understanding of modifiable and non-modifiable risk factors for osteoarthritis (OA) and post-traumatic OA (PTOA) has increased dramatically, but pathophysiological differences due to biological sex remain unclear. Gonadal hormones, namely estrogen, have been shown to modulate sex-specific responses to pain and analgesia^1–3^, and estrogen has also been established as a regulator of articular cartilage biology. Prior studies demonstrated a selective, sex-dependent response to 17β-estradiol (E2) in articular chondrocytes, which is thought to mediate cellular proliferation and matrix production. Thus, estradiol may serve a chondroprotective effect in OA.^4–6^ Concordantly, ovariectomized female mice had greater OA severity than their intact female counterparts^7^. In a human study of 196 OA patients, systemic inflammatory biomarker levels had sex-specific relationships to patient-reported knee pain: IL-1β and IL-8 levels were positively correlated to pain in male OA patients, but negatively correlated in females. IL-6 levels were negatively correlated to pain in males but positively correlated in females^8^.

Mouse models increasingly capture the diverse risk factors underlying OA development, including naturally occurring OA, use of genetic modifications, high-fat diet/obesity-induced OA, noninvasive loading-based models, and surgically-induced models^9–11^. However, for most models, comprehensive studies of sex differences and the underlying biological factors remain limited. A landmark study employing the DMM model compared OA development in intact male and female mice, orchiectomized (ORX) male mice, and ovariectomized (OVX) female mice and found that intact males exhibit greater OA severity than intact female mice^7^. Furthermore, OVX females developed worse OA than intact females, suggesting the protective effects of ovarian hormones, while males demonstrated the opposite effect as ORX mice developed less severe OA than intact males. Collectively, subsequent research using surgical mouse models has confirmed a greater overall OA/PTOA burden in males^12–15^, and it is unclear to what extent these findings may be model-dependent. Noninvasive injury models that induce joint trauma via mechanical loading offer distinct advantages for the study of joint injury, especially immediate/early manifestations. Models of repetitive joint loading^16–18^ and acute trauma^19; 20^ produce PTOA-like joint pathology and are being increasingly adopted. The tibial compression-based anterior cruciate ligament rupture (ACLR) model is simple to implement, highly reproducible, and closely recapitulates the high-energy injury conditions of human ACL injury.^9^

Despite its growing adoption, the murine ACLR model has overwhelmingly been utilized in male mice, in which it induces rapid progression of articular damage (more rapidly than surgical DMM), pronounced osteophyte formation, synovial inflammation and fibrosis, and nociceptive sensitization^19; 21–23^. The extent to which biological sex affects disease severity in the ACLR model remains largely uncharacterized, and, given recent literature on the pathogenic contribution of synovitis in PTOA^24–26^, a greater focus on inflammation and nociception is needed to comprehensively understand disease pathogenesis. The purpose of this study was to comprehensively characterize pain-, inflammation-, and joint pathology-related sex differences following noninvasive ACLR in mice (**Fig. 1**).

**Figure 1.**
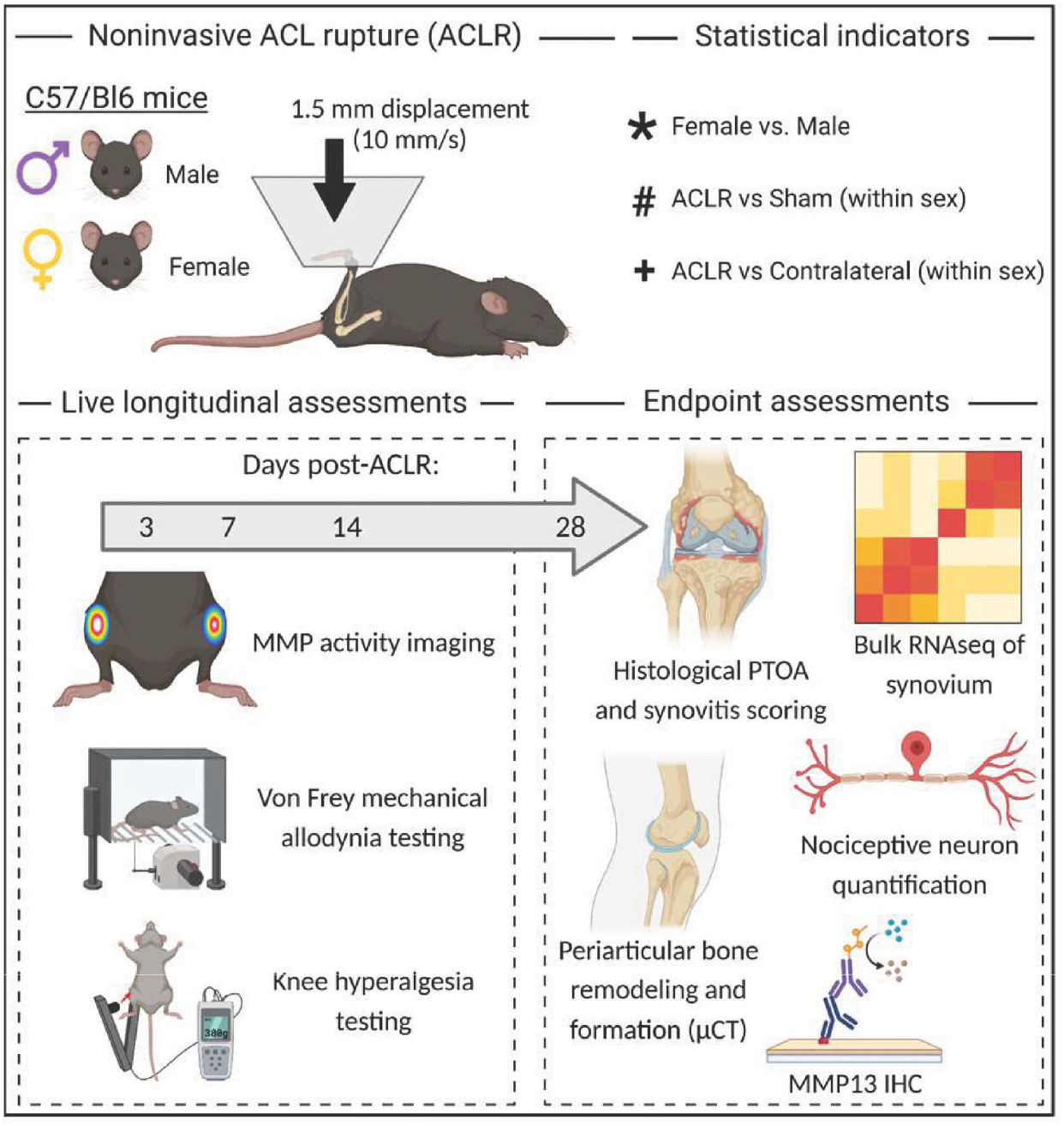
Experimental design, outcome measures, and statistical indicators.

## Methods

### Treatment Groups, Noninvasive Anterior Cruciate Ligament Rupture Model

With IACUC approval, 12-week-old male and female C57/BL6J mice (n=9 mice per group/sex/timepoint) were randomized to Sham (anesthesia and analgesia only) or noninvasive ACLR using block randomization. Within the ACLR group, mice were randomized to either 7- or 28-day endpoints, capturing pre-PTOA synovitis (7d) and established PTOA (28d)^21;27^. To power this study to evaluate sex differences, *a priori* sample size calculation was performed utilizing variance values from hyperalgesia threshold data from male mice^21^. Depending on the outcome timepoint, n=6-9 mice per sex are required to detect a 25% difference in hyperalgesia threshold, and n=9 mice/group/sex/timepoint were used. ACLR mice were subjected to a tibial compression-based ACLR protocol, as described^21^ (Supplemental Methods). Na_V_1.8-Cre;tdTomato mice^32^ (12 weeks old) also underwent ACLR to assess joint innervation (below). Mice were housed in a 12-hour light/dark facility in ventilated cages containing a maximum of 5 mice (randomized mix of Sham and ACLR) and allowed *ad libitum* activity with unrestricted access to food/water.

### Pain-related Behaviors

Knee hyperalgesia was quantified using a hand-held algesiometer (IITC Life Science, Woodland Hills, CA), as described^21^. The tip of the pressure applicator was pressed to the medial aspect of the knee joint with increasing force until a nociceptive response occurred (struggle or vocalization). An average triplicate threshold force was calculated for each limb. Mechanical allodynia in the ipsilateral hindpaw was assessed using von Frey filaments (Touch-Test Sensory Evaluators, Stoelting, Wood Dale, IL). A withdrawal response was observed using the ascending response method^28^. A response was considered positive if the animal exhibited any nocifensive behaviors. This procedure was repeated for a total of seven applications per paw per filament. The withdrawal threshold filament weight was determined when an animal responded positively to a total of 4/7 applications. Following acclimation, hyperalgesia and allodynia testing was performed longitudinally at 3d, 7d, 14d, 28d post-ACLR by a single blinded operator (RFB).

### Intravital Near Infrared (NIR) Imaging of MMP Activity

To evaluate intra-articular protease activity, we employed the protease-activatable MMPSense agent (MMPSense 680, Perkin Elmer, Waltham, MA) and intravital NIR imaging (Supplemental Methods). Briefly, MMPSense was injected into each joint and imaged 2 hrs post-injection (Pearl Impulse, LI-COR, Lincoln, Nebraska). Analysis was conducted in consistent ROIs over each joint, and raw integrated density (RID) was averaged across mice. Aggregate RID or RID normalized to the contralateral joint were compared between sexes and groups.

### Micro Computed Tomography (μCT) Imaging

Following euthanasia at 7d and 28d, skinned hindlimbs were fixed for 48 hours in 10% neutral buffered formalin, rinsed with water, and stored in 70% ethanol. One day prior to imaging, samples were rehydrated in PBS and imaged using μCT (SkyScan 1176; 8.9-μm voxel). μCT data was manually contoured to segment femoral metaphyseal trabecular bone, femoral epiphyseal trabecular bone, and medial and lateral femoral subchondral bone (SCB). Analysis was performed according to established guidelines^29^. The mean value of Sham mice within each sex were used to calculate a percent change relative to Sham, which was compared between sexes and time points (Supplemental Methods). Osteophytes were manually contoured using Dragonfly (Object Research Systems, Montreal, Canada). Total osteophyte volume was normalized to femoral length to control for skeletal size. Osteophyte mineral density and bone volume fraction were also derived, employing Otsu-based binarization.

### Structural Histology and Immunohistochemistry

Samples were paraffin processed, and 5-μm thick sagittal sections were obtained. All analyses focused on the medial joint as this compartment exhibits the greatest disease severity in the ACLR model^19^. Sections were stained with Safranin-O/Fast-Green (Saf-O) (4-6 sections/limb) and imaged at 20x (Eclipse Ni E800 with DS-Ri2 camera, Nikon, Tokyo, Japan). Blinded scoring of PTOA and synovitis^30;31^ was performed, and scores were averaged across sections within a limb and then across limbs within a group to obtain aggregate scores. Immunohistochemical staining for MMP13 was performed utilizing the Innovex Universal Animal IHC Kit and a polyclonal rabbit anti-mouse MMP13 antibody (Abcam ab39012). Briefly, sections were deparaffinized and rehydrated, and epitopes were retrieved using Uni-Trieve. After incubation with a blocking buffer, slides were incubated with primary antibody (1:75), HRP enzyme, and finally a DAB solution.

### Na_V_1.8+ Nociceptors

To visualize intra-articular nociceptors, we utilized Na_V_1.8-Cre;tdTomato mice on a C57Bl/6 background^32;33^. These mice express tdTomato in 75% of dorsal root ganglia neurons which include >90% of C-nociceptors^32^. ACLR and contralateral hindlimbs were collected from male and female Na_V_1.8-Cre;tdTomato mice (n=3) following transcardial perfusion with 4% PFA at 28d post-ACLR. Twenty-μm thick coronal sections were collected from the mid-joint area and imaged by confocal microscopy as described^34^. Na_V_1.8+ signal in the medial synovium and the number of Na_V_1.8+ SCB channels were quantified in a blinded fashion using ImageJ (Supplemental Methods).

### RNA Sequencing of Synovium and Bioinformatic Analyses

Whole synovium, inclusive of Hoffa’s fat pad, was microdissected from male and female Sham, contralateral, and ACLR knee joints (n=6/sex/group), as previously described^21^. A standard workflow for RNA isolation, quality control, bulk RNA-sequencing, and bioinformatic analyses of differentially-expressed genes (DEGs) and differentially-regulated pathways was used to assess sex-independent and sex-dependent perturbation to the synovial transcriptome following ACLR (Supplemental Methods).

### Statistical Analyses

SPSS (v27, IBM, Armonk, NY) and Prism 9.0 (Graphpad, San Diego, CA) were used for statistical analyses. Four-way linear mixed-effects models with Sidak post-hoc correction compared longitudinally-measured, repeated/live, continuous outcomes in the Sham and ACLR groups (limb and time point as within-subject factors, and group and sex as between-subject factors). Three-way linear mixed-effects models with Sidak post-hoc tests compared continuous outcomes within the ACLR group whenever both limbs were analyzed in separate groups/sexes of mice at different time points (limb as a within-subject factor, and time point and sex as a between-subject factor). These outcomes were compared between either limb in the ACLR group to the single independent sex-matched Sham group using one-way ANOVA with Dunnett’s post-hoc test (Sham as the control group). Histological scores and non-normally distributed data were analyzed between matched limbs within male and female ACLR mice using Wilcoxon Signed Rank tests with Sidak family-wise *P* value correction. Scores and non-normally distributed data were compared between time points within the ACLR group (by sex) or between sex-matched Sham and ACLR mice using Kruskal-Wallis tests with Dunn’s post-hoc tests. Sparse partial least squares-discriminant analysis (sPLS-DA) was used to perform dimensionality reduction of raw, non-normalized μCT data, and variable importance in projection (VIP) coefficients were calculated for the components describing sex-based and injury-induced variance. Corrected *P* values below 0.05 were considered significant throughout.

## Results

### ACLR induces progressive joint damage and synovitis in both sexes, with greater severity in male mice

Male and female mice exhibited typical histological features of PTOA following ACLR: cartilage erosion and fibrillation, loss of cartilage proteoglycan content, meniscal damage, osteophyte formation, and SCB thickening (**Fig. 2A**). In both sexes, knee joints exhibited greater PTOA scores 7d and 28d after ACLR relative to their contralateral joints (**Fig. 2B**). Only male ACLR joints had greater PTOA scores relative to male Sham at 7d, and both sexes exhibited greater PTOA scores relative to their respective Sham at 28d. At both timepoints, males had significantly greater PTOA scores compared to females (**Fig. 2A-B)**, driven most notably by greater structural damage (loss of noncalcified cartilage), proteoglycan loss in noncalcified cartilage, and osteophyte size compared to females (**Suppl. Fig. S2A).** We observed no notable joint damage or increased PTOA scores in contralateral joints (**Fig. 2B-C**).

**Figure 2.**
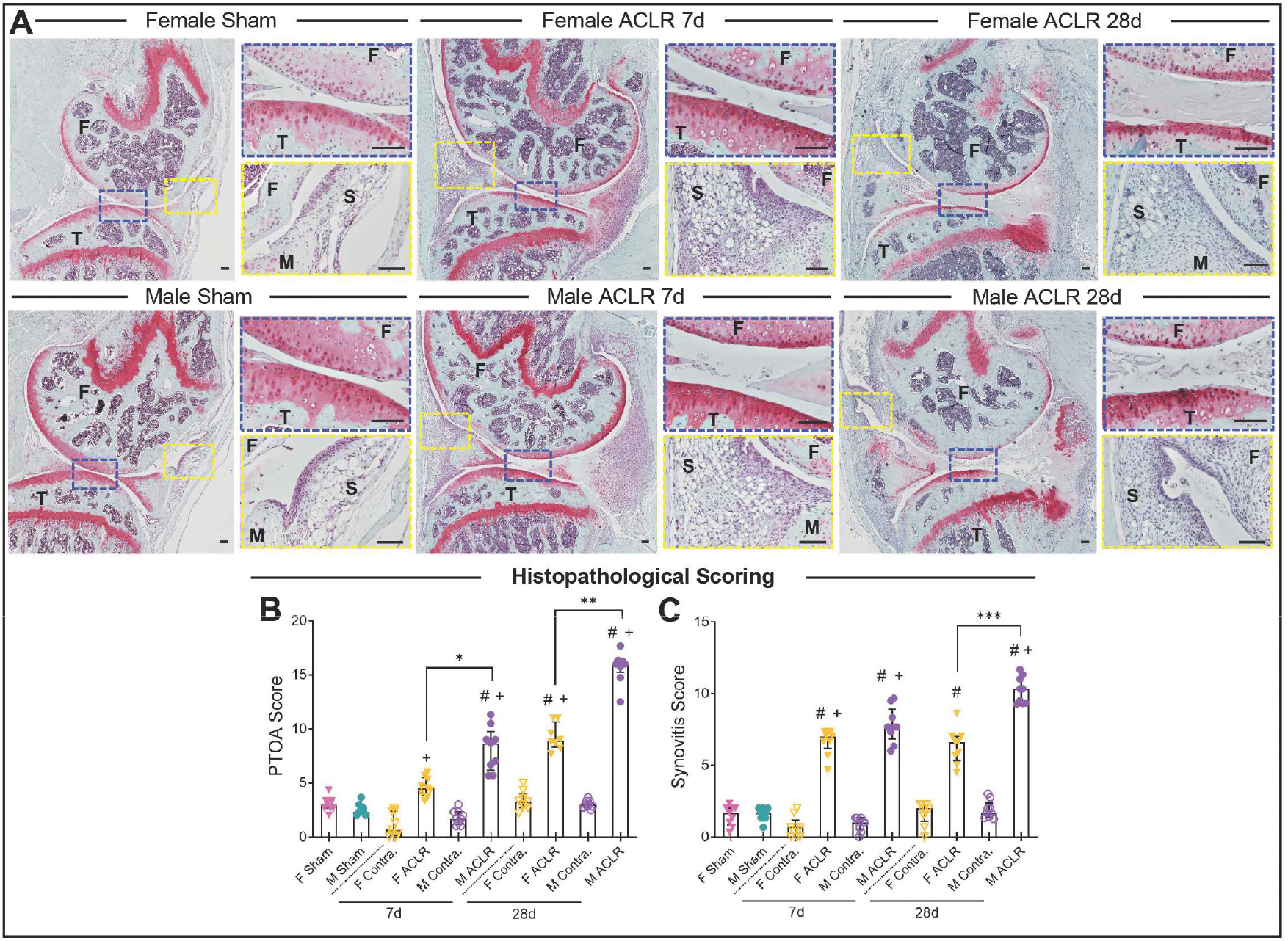
Male mice exhibit greater PTOA and synovitis severity. **A).** Representative sagittal histological sections of the medial joint compartment stained with Safranin-O/Fast-Green in male and female Sham, ACLR 7d, and ACLR 28d mice. Scale bar 100 μm. **B-C).** Histopathological scoring of PTOA (**B**) and synovitis (**C**) in Sham, Contralateral (Contra) and ACLR limbs. n=8-9 limbs per group. Error bars represent interquartile range. * P < .05 between males and females. # P < .05 between ACLR and Sham within each sex. + P < .05 between ACLR and Contralateral within each sex.

Histopathological scoring further revealed robust synovitis in both males and females by 7d, with both sexes exhibiting greater synovitis scores relative to their contralateral joints and Sham (**Fig. 2C**). The synovitis scores of male mice continued to increase between 7d and 28d, but this was not observed in females, and at 28d, males exhibited significantly greater synovitis scores than females. This was marked by greater lining hyperplasia, more severe subsynovial inflammatory infiltrate (**Suppl. Fig. S2B)**, and worse fibrosis in males, whereas female synovia exhibited greater maintenance of normal areolar synovium (**Fig. 2A**). Taken together, these results indicate that male mice exhibit worse progressive PTOA severity and synovitis following ACLR relative to female mice.

### Male mice exhibit greater intra-articular MMP activity

Both sexes exhibited elevated normalized MMP activity (ACLR/Contra ratio) following injury compared to Sham, observed as early as 3d, with sustained elevated MMP activity until 28d (**Fig. 3A-B**). Only male mice exhibited progressive increases in MMP activity up to 28d, at which point males had significantly greater normalized MMP activity relative to females (**Fig. 3B**). Contralateral joints demonstrated similar MMP activity as Sham mice. To spatially contextualize the cell and tissue types underpinning injury-induced MMP activity, we performed immunohistochemical staining against MMP13, an MMP member relevant to cartilage and bone erosion, chondrocyte hypertrophy, and osteophyte formation. Qualitative evaluation demonstrated MMP13+ cells in the synovial sublining layer, but not synovial lining, in articular chondrocytes, meniscal fibrochondrocytes, and osteophyte-resident hypertrophic chondrocytes (**Fig. 3C**). These findings indicate that joint injury induces greater longitudinal MMP activity in male mice, likely contributing to the greater cartilage damage observed in males.

**Figure 3.**
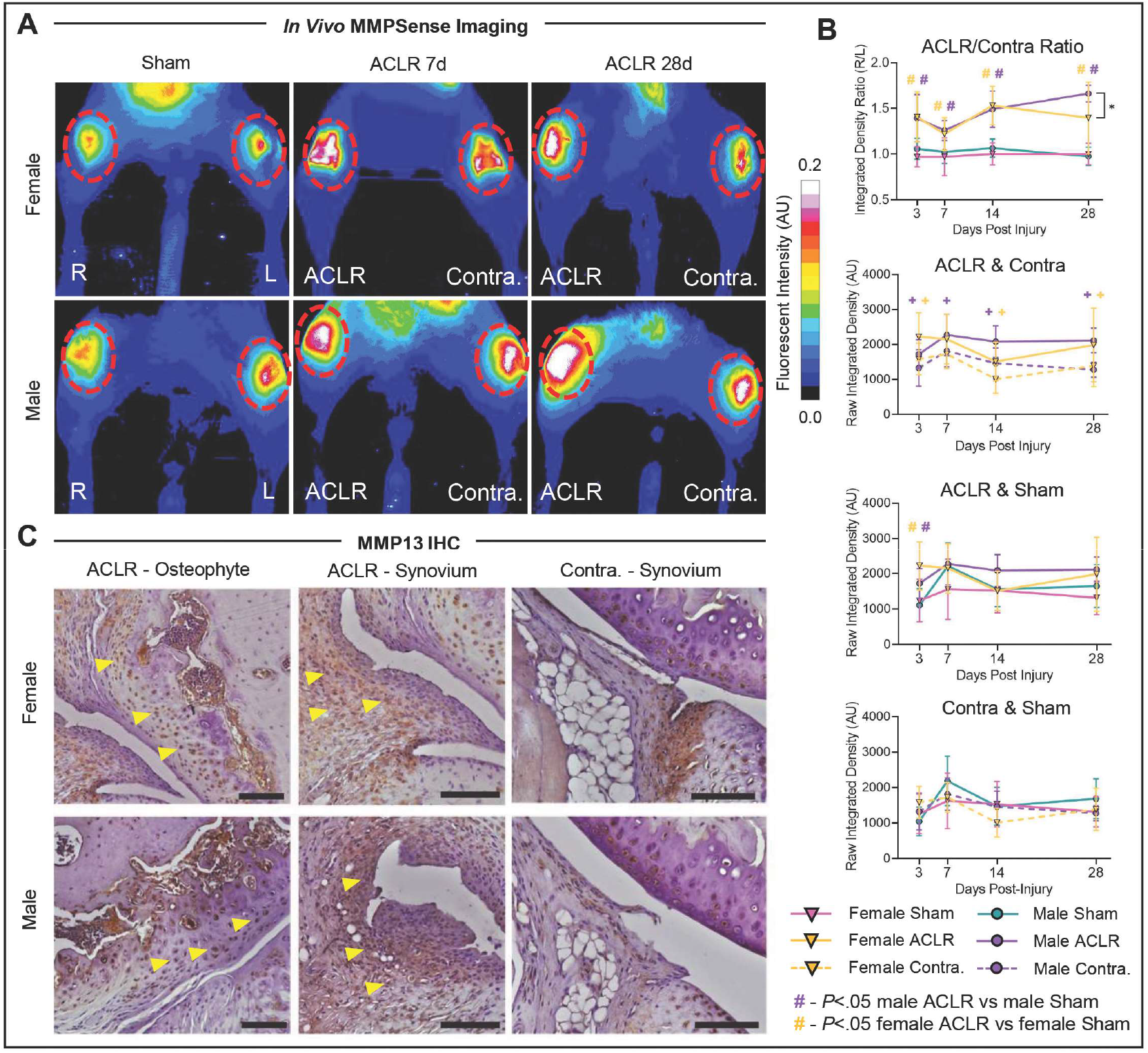
Male mice exhibit greater injury-induced intra-articular MMP activity. **A**). Representative intravital fluorescence imaging heatmap of MMP activity. **B).** Longitudinal MMPsense quantification, expressed as raw integrated fluorescence density or integrated density ratio (right/left) in Sham, ACLR, and Contralateral joints. n=6-9 limbs per group. Error bars represent 95% confidence interval. * P < .05 between males and females. # P < .05 between ACLR and Sham within each sex. + P < .05 between ACLR and Contralateral within each sex. **C).** MMP13 immunostaining in male and female ACLR and contralateral joints. Yellow arrowheads indicate MMP13+ cells. Scale bar is 100 μm.

### Males and females exhibit differential injury-induced bone remodeling, and males exhibit greater osteophyte formation

Males and females exhibited markedly different trabecular and SCB remodeling post-ACLR, with complex compartment- and parameter-dependent patterns (**Fig. 4A-D**). Females exhibited notable injury-induced trabecular bone loss, mineral density decreases, and trabecular thinning in the metaphysis and epiphysis, whereas males had little to no catabolic remodeling (**Fig. 4A-B, Suppl. Fig. S3**). Further, females, but not males, exhibited medial and lateral SCB thickness loss at 7d (**Fig. 4C-D**). Only males exhibited increases in medial SCB thickness and mineral density, indicative of SCB sclerosis, at 28d (**Fig. 4C**), consistent with greater observed histological PTOA severity, and both sexes showed loss of lateral SCB mineral density at 28d (**Fig. 4D**).

**Figure 4.**
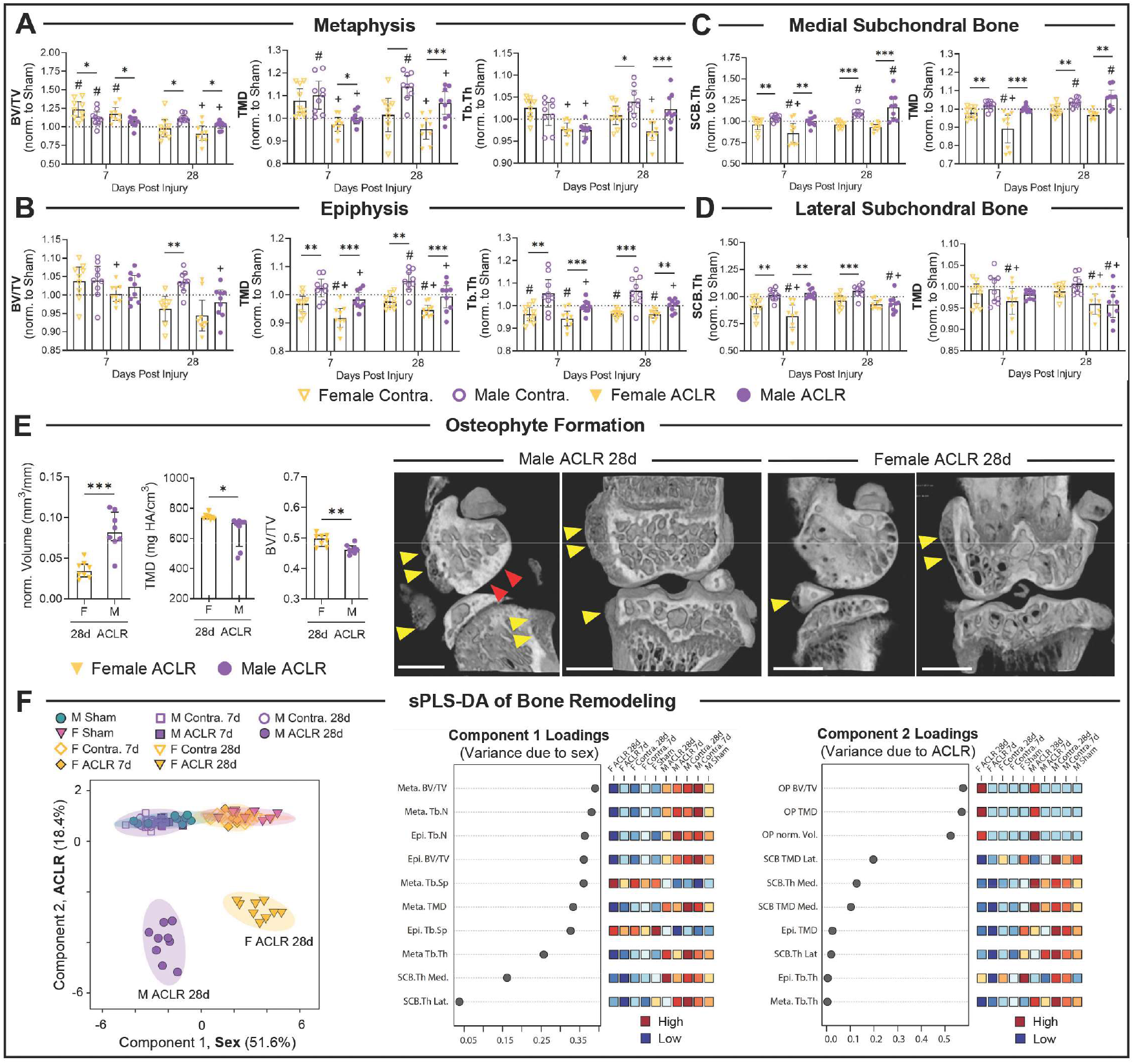
Male and female mice exhibit differential bone remodeling patterns following joint injury. **A-D).** Morphometric parameters derived from μCT in metaphyseal trabecular bone (**A**), epiphyseal trabecular bone (**B**), medial subchondral bone (**C**), and lateral subchondral bone (**D**). **E).** Osteophyte formation morphometric parameters (left) and representative 3-dimensional reconstructions (right). Yellow arrowheads point towards osteophytes in male and female ACLR 28d joints. Red arrowheads point towards medial subchondral bone sclerosis. Scale bar is 1 mm. **A-E).** n=8-9 limbs per group. Error bars represent 95% confidence interval. * P < .05 between males and females. # P < .05 between ACLR and Sham within each sex. + P < .05 between ACLR and Contralateral within each sex. **F).** Scores plot (left) and variable importance in projection loadings from component 1 (middle) and 2 (right) from sparse partial least squares discriminant analysis (sPLS-DA) of μCT-derived trabecular, subchondral, and osteophyte parameters. n=8-9 limbs per group.

Osteophyte formation was observed in similar patterns in both sexes, most pronounced in the medial compartment, and not evident until 28d (**Fig. 4E**). Normalized osteophyte (OP) volume was greater in ACLR males than females at 28d. Conversely, OP maturity, as determined by TMD and BV/TV, was slightly greater in ACLR females than males (**Fig. 4E**).

To reduce the dimensionality of these complex remodeling patterns, we performed sPLS-DA. The resultant score plot demonstrates sex-based variance on Component 1 and injury-based separation on Component 2 (**Fig. 4F**). Strikingly, despite our observations of injury-induced remodeling at 7d (**Fig. 4A-D**), we observed no separation between sham, contralateral, and 7d ACLR limbs in either sex (**Fig. 4F**), whereas male and female 28d ACLR limbs exhibited clear separation. VIP coefficients showed that epiphyseal and metaphyseal trabecular parameters dictated sex-based variance, whereas osteophyte and SCB parameters were most contributory to injury-induced variance (**Fig. 4F**). Taken together, males exhibit greater injury-induced sclerotic remodeling and osteophyte formation, consistent with greater PTOA severity observed in males, whereas females have notable catabolic trabecular and SCB responses to injury.

### Both sexes show comparable nociceptor sprouting after ACLR, but males develop more knee hyperalgesia and mechanical allodynia

Both sexes showed a significant reduction in knee and hind paw withdrawal thresholds between 3-28d post-ACLR, compared to sex-matched Shams (**Fig. 5A-B**) and contralateral limbs (**Suppl. Fig. S4**). These measures began to diverge between the sexes at 14d, and significant differences in both knee hyperalgesia and mechanical allodynia were observed between males and females by 28d (**Fig. 5A-B**). We also observed sensitization in the contralateral limb of both sexes, with significant differences in knee withdrawal threshold at 7d, 14d, and 28d in both sexes compared to Shams (**Fig. 5A-B**). Mechanical allodynia changes were also detectable but more subtle in contralateral joints, and males had a significantly lower contralateral hindpaw withdrawal threshold compared to females at 28d. To contextualize pain behavior with histological evidence of nociceptor sprouting, we used Na_V_1.8-Cre; tdTomato mice, which express tdTomato in nociceptive neurons. At 28d, we observed tdTomato+ SCB channels, most notably in the medial compartment, and extensive tdTomato+ fibers in the medial synovium of both male and female mice (**Fig. 5C**). Contralateral joints of either sex did not exhibit these features. Male and female ACLR joints had significantly greater tdTomato+ synovial surface area (**Fig. 5D**) and number of SCB channels (**Fig. 5E**) compared to contralateral joints, but no differences were observed between males and females. Taken together, both males and females exhibit injury-induced knee hyperalgesia and mechanical allodynia, with greater severity in males by 28d. These changes are associated with the sprouting of nociceptors into SCB and synovium, which was observed at comparable levels between the sexes.

**Figure 5.**
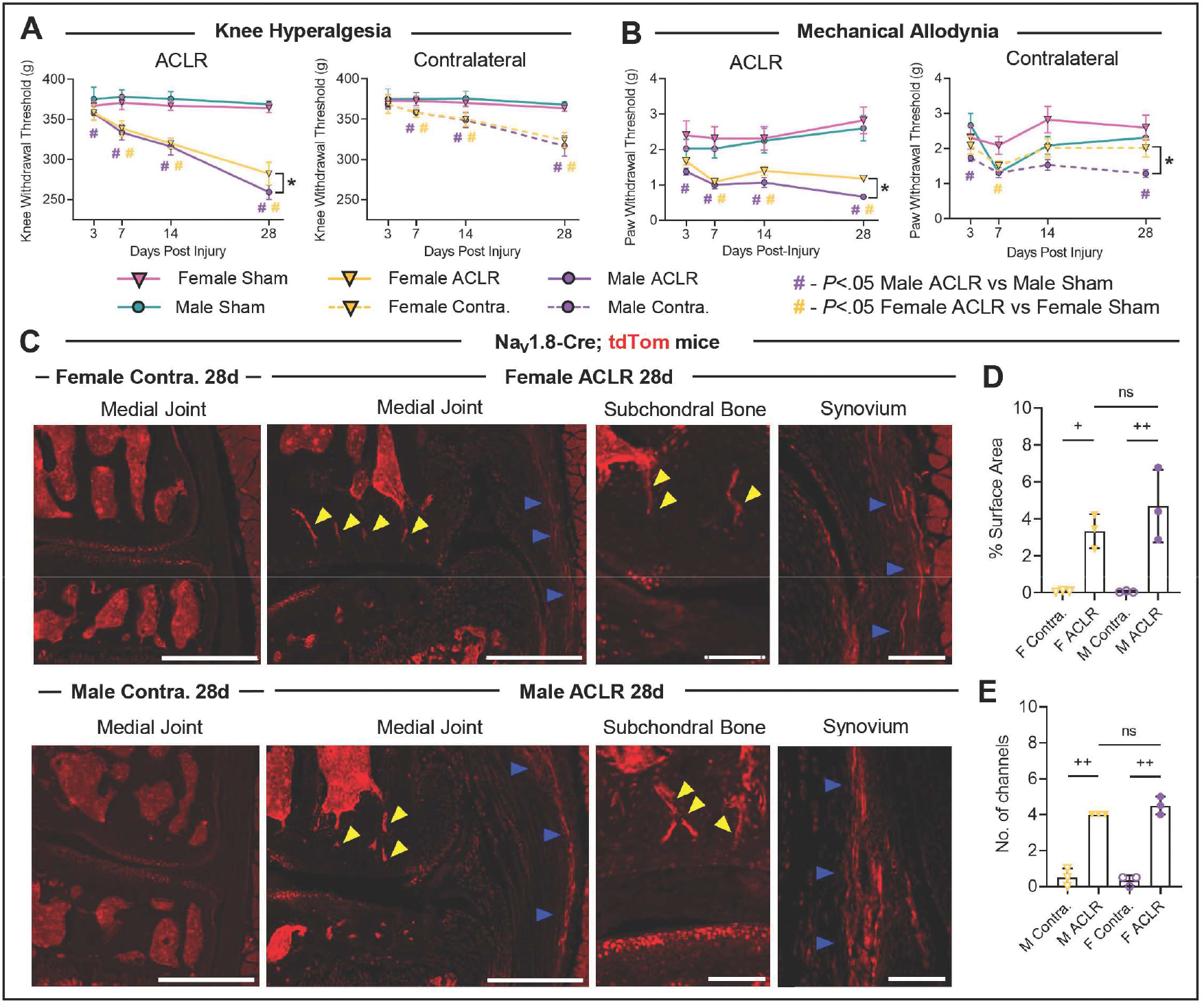
Male mice exhibit greater injury-induced knee hyperalgesia and mechanical allodynia. **A-B).** Longitudinal assessment of knee hyperalgesia (**A**) and mechanical allodynia (**B**) in ACLR and Contralateral joints, compared to sex-matched Sham controls. n=9 mice per group. Error bars represent 95% confidence interval. * P < .05 between males and females. # P < .05 between ACLR and Sham within each sex. **C).** Fluorescent microscopy of coronal joint sections of Na_V_ 1.8-Cre; tdTom mice in male and female ACLR 28d and respective Contralateral joints. Yellow arrowheads indicate tdTom+ subchondral bone channels. Blue arrowheads indicate tdTom+ fibers in synovium. Large scale bar is 500 μm. Small scale bar is 100 μm. **D-E)**. Quantification of synovial tdTom+ fibers by proportion of total surface area (**D**) and tdTom+ subchondral bone channels by total number of channels per section (**E**) in 28d ACLR and Contralateral joints. n=3 per group. + P < .05 between ACLR and Contralateral within each sex. ++ P < 0.001 between ACLR and Contralateral within each sex. ns – not significant.

### RNA-Seq of synovium reveals inflammatory and fibrosis-related pathway activation following ACLR

Our findings indicating greater MMP activity, synovitis, and pain in males prompted us to undertake bulk RNA-seq of synovium to identify the molecular changes underpinning these differences. ACLR induced massive perturbation to the synovial transcriptome, with greater than 10,000 DEGs (P_adj_<0.05) at 7d and 28d relative to Sham (**Fig. 6A**). DEGs for all comparisons are in Supplemental Table 1. GO pathway analysis revealed activation of the inflammatory and immune response, with both myeloid and lymphocyte-relevant pathways, enriched extracellular matrix remodeling processes indicative of synovial fibrosis and endochondral tissue formation, and enhanced neuroangiogenesis (**Fig. 6B**). Consistent with the fibrosis-associated loss of the fat pad, observed histologically in our mice (**Fig. 2A**) and known to occur in human OA^35–37^, we observed downregulation of adipogenic pathways such as “brown fat cell differentiation” (**Fig. 6B**). Wnt/β-Catenin signaling, integrin-mediated signaling, and ERK1/2 signaling were activated at both 7d and 28d, and Smoothened/Hedgehog signaling was enhanced at 28d (**Fig. 6B).** We observed very few DEGs and no significantly regulated pathways between Sham and Contralateral synovia at either timepoint (**Suppl. Fig. S5A-B, Suppl. Table 1**), and the majority of DEGs were overlapping between the Sham vs ACLR and Contralateral vs ACLR comparisons (**Suppl. Fig. S5B**). Next, we assessed transcriptomic changes associated with PTOA progression, comparing ACLR 7d to ACLR 28d (**Fig. 6C**). Relative to Sham, injury-induced DEGs were largely overlapping between 7d and 28d (**Fig. 6D**). Pathway analysis of DEGs between ACLR 7d and 28d revealed downregulation of immune and inflammatory response-related pathways at 28d (**Fig. 6E**), indicative of partial resolution yet chronic maintenance of synovitis. ECM-related pathways indicative of fibrosis (Notch signaling) and endochondral tissue formation (Cellular response to BMP stimulus) were enriched at 28d relative to 7d, and cell cycle and cell division-related pathways were downregulated at 28d (**Fig. 6E**). The senescence marker *p16/Cdkn2a* was upregulated at 28d and associated with downregulation of the “cell division” GO pathway (**Fig. 6F**), suggesting the enrichment of senescent cells in ACLR 28d synovium.

**Figure 6.**
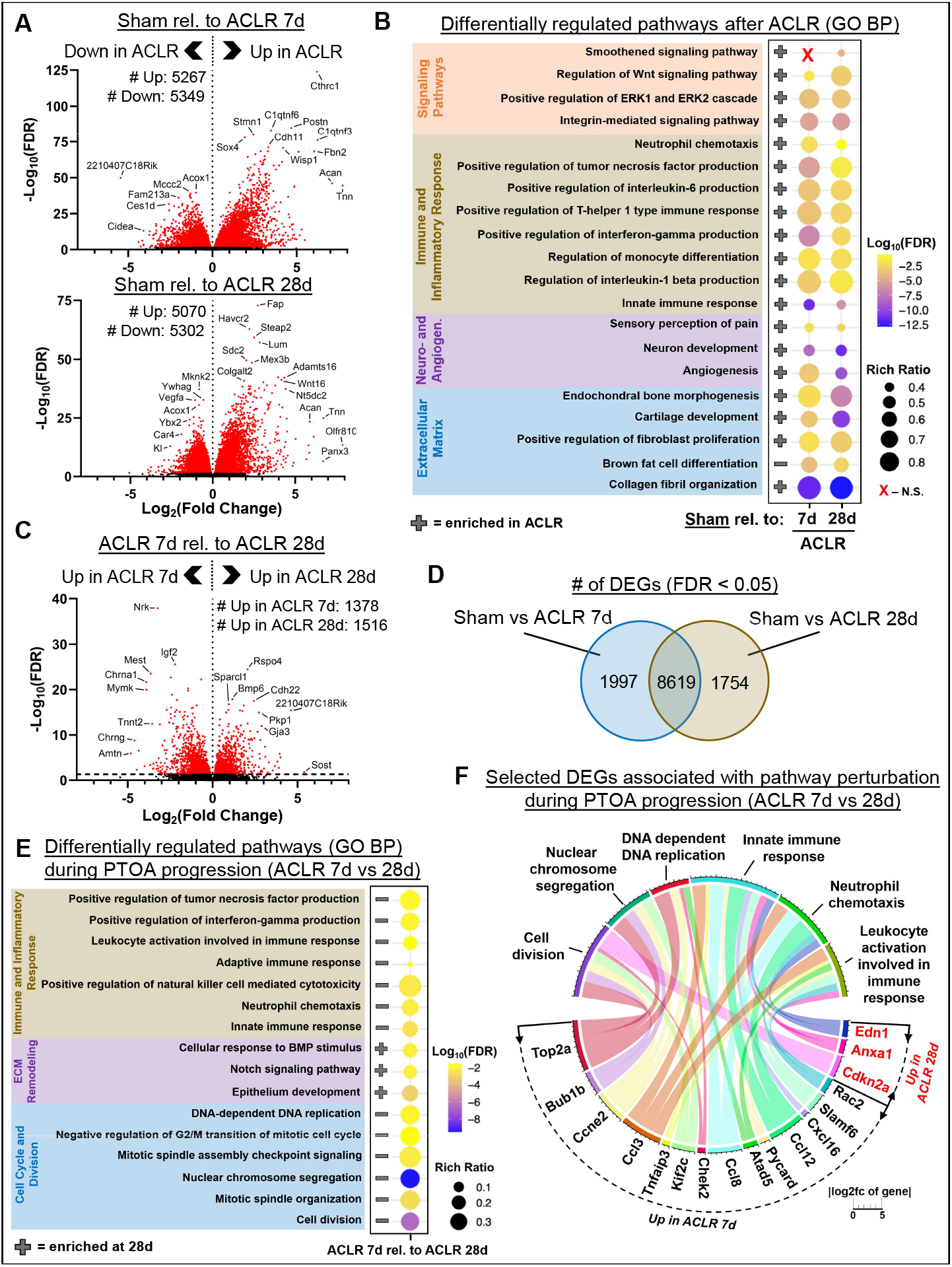
RNA-seq reveals pathway activation signature of synovium following joint injury. **A).** Volcano plots showing differentially expressed genes (DEGs) in ACLR 7d (top) and ACLR 28d (bottom) relative to Sham. **B).** Differentially activated pathways derived from the DEGs from (**A**) using the Gene Ontology (GO) “Biological Process” annotation. The direction of activation is indicated by +/-, with + indicating enrichment in ACLR relative to Sham. **C).** Volcano plots showing DEGs between ACLR 7d and ACLR 28d. **D).** Venn diagram illustrating overlapping DEGs between Sham vs ACLR 7d and Sham vs ACLR 28d. **E).** Differentially activated pathways during PTOA progression, derived from the DEGs from (**C**) using the GO:BP annotation. The direction of activation is indicated by +/-, with + indicating enrichment in ACLR 28d relative to ACLR 7d. **F)**. Chord plot of DEGs associated with the regulation of GO:BP pathways from (**E**). Genes in black font are over-expressed in ACLR 7d, and genes in red font are over-expressed in ACLR 28d. **A-F).** n=11-12 synovia per condition (n=5-6 male and female per condition).

### Sex differences in the synovial transcriptome are marked by greater maintenance of pro-fibrotic and pro-inflammatory signaling, neuroangiogenesis, and mineralization in male mice

Comparing the synovial transcriptome of male and female ACLR mice directly, we unexpectedly found only 80 DEGs unique to 7d, 265 DEGs unique to 28d, and 44 DEGs overlapping both timepoints (**Fig. 7A-B**). At 7d, GO analysis revealed no differentially perturbed pathways between males and females, indicating that the early injury-induced synovial response is similar between the sexes. At 28d, pathway analysis demonstrated enriched pathways in male synovia indicative of fibrosis, neuroangiogenesis, and mineralization (**Fig. 7C**). Gene markers associated with fibroblast activation and collagen production, *e.g. Tnn, Cdh11, Col1a1, Col3a1, Lox*, and markers of neuroangiogenesis, *e.g. Sema3c, Igf1, Angptl4, Nrxn1, Ncan, Postn* were more highly expressed in male synovia compared to female synovia at 28d (**Fig. 7C, Suppl. Table 1**). Given these divergent transcriptomes at 28d, we assessed whether males and females exhibit differential disease progression. We found 2706 DEGs unique to female PTOA progression (**Fig. 7D**), representing pathways indicative of the resolution of inflammatory and fibrotic processes (**Fig. 7E**). The PCA scores plot further corroborates resolution of injury-induced gene expression only in female ACLR 28d, whereas male ACLR 28d synovia cluster together with 7d ACLR synovia of both sexes (**Suppl. Fig. S5C-D**). We further found enhanced pathways related to fatty acid metabolism and responses to insulin, indicating differences in adipose metabolism between the sexes. Taken together, these findings show that while both sexes exhibit activation of inflammatory, immune, and fibrotic processes following joint injury, female mice exhibit enhanced resolution of these processes whereas males maintain transcriptional programs promoting chronic inflammation and fibrosis.

**Figure 7.**
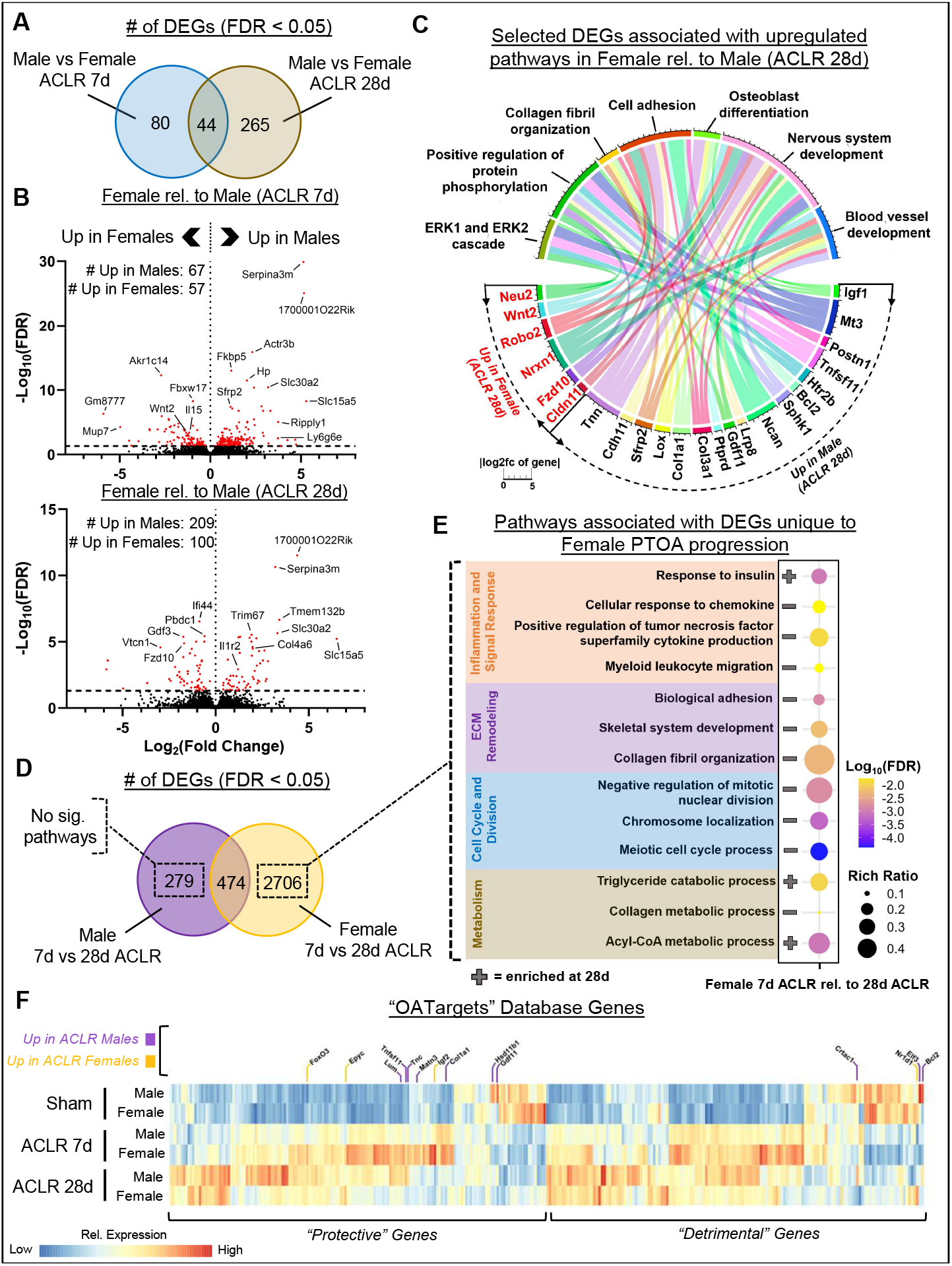
Female synovia exhibit inflammatory resolution, whereas male synovia have persistent pro-inflammatory and pro-fibrotic gene expression. **A).** Venn diagram showing overlapping DEGs between male vs female ACLR at 7d and male vs female ACLR at 28d. **B).** Volcano plots showing DEGs in male vs female ACLR at 7d (top) and 28d (bottom). **C**). Chord plot of DEGs associated with the regulation of GO:BP pathways from male vs female ACLR at 28d. Genes in black font are over-expressed in male ACLR 28d, and genes in red font are over-expressed in female ACLR 28d. All GO:BP pathways shown are upregulated in males relative to females. **D).** Venn diagram showing overlapping DEGs between 7d vs 28d ACLR in males and 7d vs 28d ACLR in females. **E).** Differentially activated pathways derived from the DEGs from female 7d vs 28d ACLR using GO:BP annotation. The direction of activation is indicated by +/-, with + indicating enrichment in female ACLR 28d relative female ACLR 7d. **F).** Heatmap of genes in the OATargets “protective” and “detrimental” categories across male and female Sham, ACLR 7d, and ACLR 28d synovia. Select genes with differential expression between males and females are listed, with the color corresponding to direction of expression. **A-F).** n=5-6 synovia per condition.

We lastly analyzed how known OA-relevant genes were altered in our model by mapping the “OATargets” database (459 genes)^38^ to our results (**Fig. 7F**). We observed massive changes in both the “protective” and “detrimental” gene categories following ACLR, with clear sex-dependent expression profiles. Similar patterns were observed in the “ambiguous” gene category (**Suppl. Fig. S5E**), representing genes without a consensus effect on OA progression^38^. No clear pattern was observed regarding one sex being over/underrepresented in either the “protective” or “detrimental” category.

## Discussion

We uncovered pronounced sex differences in mechanical sensitization, tissue damage/remodeling, synovitis, and the synovial transcriptome in the mouse ACLR model. While both sexes exhibited characteristic histological features of PTOA, male mice exhibited greater pain-related behaviors, synovitis, and structural damage than female mice up to 28d. Our findings extend prior research in models of spontaneous and induced OA^14;39–41^ by demonstrating that after ACLR, greater overall disease severity in male mice – including joint pathology and associated pain behaviors – is associated with a divergence in the synovial transcriptome between the sexes, marked by the absence of resolution of synovial inflammation and fibrosis in males.

The contribution of synovitis to OA pathology^24–26^ and the association between synovitis and pain are increasingly recognized^42^. We present evidence that worse pathology in male mice is associated with greater synovial gene expression relevant to fibrosis, inflammation, neuroangiogenesis, and mineralization, whereas females exhibited partial resolution of these processes. Our RNAseq results should be interpreted in light of the recent publication of a single-cell RNAseq atlas of mouse sham, ACLR 7d, and ACLR 28d synovium by our group^27^, which utilized pooled male and female samples to demonstrate the dramatic enrichment and diversification of synovial fibroblasts and macrophages following injury. In the present study, the resolution signature observed in female ACLR mice, but not males, demonstrating downregulated myeloid migration, cellular responses to chemokines, and collagen fibril organization is consistent with either decreases in synovial fibroblast and macrophage numbers, downregulation of their cellular activity, or both. Studies of human synovium have previously linked patient-reported pain to synovitis, synovial gene expression, and cellular composition. Scanzello *et al*. showed that in patients undergoing arthroscopic meniscectomy, histologically-diagnosed synovitis was associated with worse preoperative pain and function scores, and patients with inflammation had increased synovial gene expression of *IL-8, CCL5, CCR7*, and *CCL19^43^*. Nanus *et al*. showed that synovial tissue from painful sites in early OA patients had enriched activation of pro-fibrotic (e.g. *TGFB1, FZD7*), pro-inflammatory (e.g. *TRAF2*, a regulator of NF-κB), and neurotrophic signaling (e.g. *NRN1, CREB3)^44^*. scRNA-seq demonstrated that painful sites had greater abundances of a fibroblast subset enriched in IGF-1 signaling, corroborating our data that males had greater synovial *Igf1* expression (**Fig. 7C**). Relevant to male mice exhibiting persistent activation of the “cellular response to chemokine” pathway, numerous studies have established the direct catabolic effect of chemokines on chondrocytes, their role in recruiting pro-inflammatory immune cells, and their regulation of mechanical sensitivity and nociceptor sprouting in OA^24; 45; 46^. Thus, our findings suggest that PTOA in murine males involves persistent activation of pathologic signaling in synovium, which likely promote worse nociceptive and structural changes.

Multiple results confirmed greater joint damage in male mice. Our data extend prior evidence of worse cartilage pathology in male mice by associating it with greater intra-articular MMP activity, a process relevant to proteolytic cartilage matrix loss and pathological chondrocyte hypertrophy observed in SCB sclerosis and osteophyte formation. Bone-related indicators of sex-dependent PTOA severity were primarily reflected in osteophyte formation, which was markedly higher in males, and SCB sclerosis, which was only observed in males. Metaphyseal and epiphyseal trabecular bone exhibited pronounced sex differences but no appreciable injury-induced remodeling pattern. Satkunananthan *et al*. observed no sex differences in osteophyte or histological OA severity 8-weeks following ACLR in C57Bl/6 mice^47^, but this study did not evaluate nociceptive changes or earlier timepoints. Given studies in other mouse OA models demonstrating greater disease severity in males at established/late-stage disease^7; 14; 39–41^, our findings are likely not isolated to the early post-injury period up to 28d.

ACLR caused immediate ipsilateral knee hyperalgesia and mechanical allodynia in both sexes. Mechanical allodynia was sustained over 28d, while knee hyperalgesia showed progressive worsening. At the time of established PTOA, males exhibited worse knee hyperalgesia and mechanical allodynia, consistent with the worse joint damage and synovitis we observed. Several studies have reported pain-related behaviors associated with experimental OA in male mice, but very few have directly compared progression of OA and pain between males and females. Temp *et al*. showed that male C57Bl/6 mice underwent a full remission period in mechanical allodynia between weeks 1-3 following surgical medial meniscal transection (MMT), whereas females exhibited no remission but sustained allodynia^14^. They found no differences in mechanical allodynia in the chronic phase. von Loga *et al*. found similar pain-like behavior between the sexes after DMM, measured by hindlimb weight bearing, but females had greater neurotrophic gene expression in whole-joint samples (e.g. *Gdnf, Nrtn, Ntf3, Ntf5*)^15^. Ishihara *et al*. reported a large decrease (30-40%) in knee withdrawal threshold 2 weeks after DMM surgery in male C57Bl/6, followed by a linear recovery of hyperalgesia up to week 16^46^. In female C57Bl/6 mice induced by partial meniscectomy, Knights *et al*. observed an early knee hyperalgesia phase between day 7 and 14, a recovery to baseline between day 21 and 28, followed by a chronic hyperalgesia phase from day 35 until the study end at day 84, whereas mechanical allodynia was not elevated beyond the level of sham-treated mice until day 56 and beyond^48^. These studies found allodynia and hyperalgesia patterns strikingly different to the pain progression data in our study, strongly indicating that longitudinal pain responses in the ACLR model are distinct from surgical models.

Our transcriptomic results confirm greater neurotrophic gene expression in male synovia, and although we observed worse long-term pain in males, we did not find greater tissue densities of NaV1.8-tdTomato+ nociceptors. These results are limited by the small sample size of the Na_V_1.8-tdTomato cohorts (n=3) and the use of separate C57Bl/6 and Na_V_1.8-tdTomato cohorts. Future work is necessary to comprehensively map the nociceptive innervation of the knee joint in various OA models to understand whether worse pain signatures are associated with greater tissue densities of nerve fibers. Sex differences studies in the afferent signal-processing entities, i.e. dorsal root ganglion, will further contextualize model-dependent pain responses and joint pathology.

In conclusion, we demonstrated marked sex-based differences in both joint pathology and pain-related behaviors in the murine ACLR model, and a synovial inflammatory resolution signature underpinned milder disease and lesser pain in female mice. These results establish the ACLR model as a useful tool to study trauma-induced pain behavior relevant to PTOA and support the role of chronic synovitis, marked by immune cell enrichment, fibrosis, and mineralization, as a driver of pathology and pain.

## Supplemental Figure Legends

**Suppl. Figure S1 – A).** Body weight and descriptive mechanical properties derived during the anterior cruciate ligament rupture (ACLR) procedure between male and female mice. **B)**. Linear correlations between weight and failure load (left) or failure displacement (right). n=18 per sex. **** indicates P < 0.0001.

**Suppl. Figure S2** – Histopathological PTOA (**A**) and synovitis (**B**) subscores. n=8-9 per group. Error bars represent interquartile range.

**Suppl. Figure S3** – μCT supplement. n=8-9 limbs per group. Error bars represent 95% confidence interval. * P < .05 between males and females. # P < .05 between ACLR and Sham within each sex. + P < .05 between ACLR and Contralateral within each sex.

**Suppl. Figure S4** – Comparison of ACLR and Contralateral within each sex in knee withdrawal threshold from knee hyperalgesia testing (**A**) and paw withdrawal threshold from mechanical allodynia testing (**B**). n=9 per group. * P < .05 between male and female. + P < .05 between ACLR and Contralateral within each sex.

**Suppl. Figure S5** – **A).** Volcano plots showing DEGs (red) between Sham vs Contra 7d and Sham vs Contra 28d comparisons. **B).** Venn diagrams showing overlap of DEGs between Sham vs ACLR and Contralateral (Contra) vs ACLR at 7d (top) and 28d (bottom). **C-D).** Principal component analysis scores plot (**C**) and top/bottom loadings (**D**) for component 1 (top) and component 2 (bottom) from bulk RNA-sequencing of synovium. n=5-6 per condition. **E).** Heatmap of genes in the OATargets “ambiguous” category across male and female Sham, ACLR 7d, and ACLR 28d synovia. Select genes with differential expression between males and females are listed, with the color corresponding to direction of expression.

## Supplemental Table Figure Legends

**Suppl. Table 1:** Differentially expressed gene (DEG) lists from all comparisons across sexes, groups, and timepoints from bulk RNA-seq of synovium.

