## Supplementary material for "Sexual dimorphism of the synovial transcriptome underpins greater PTOA disease severity in male mice following joint injury": Supp. Methods

**Supplemental Methods:**

*Noninvasive Anterior Cruciate Ligament Rupture Model*

Injuries were conducted as previously described^25^. Mice were anesthetized with 2% inhaled isofluorane and immobilized on a custom fixture on a mechanical testing system (Electroforce 3300AT, TA Instruments, New Castle, DE). The right knee was flexed to 100^o^ and secured in a trough to restrict medial-lateral motion. The hindpaw was secured in 30^o^ dorsiflexion. After preloading and conditioning, a rapid displacement of 1.5 mm (10 mm/s) was applied via the hindpaw, causing tibial compression, anterior tibial subluxation, and ACL rupture. Immediately after ACLR, mice received a single dose of subcutaneous carprofen (5 mg/kg). Complete ACL rupture was confirmed via an anterior drawer test. Failure/rupture load, failure displacement, and linear elongation stiffness were derived from mechanical data sampled during the injury procedure (100 Hz) and compared between male and female mice (**Suppl. Fig. S1**). Body weight was expectedly higher in male mice (P < .0001). Male mice had marginally higher failure load (P = .086) and failure displacement (P = .059), but no difference in linear elongation stiffness was observed between the sexes (P = .974) (**Suppl. Fig. S1A**). Body weight weakly correlated to failure displacement (r^2^ = .122, P = .037) but not to failure load (r^2^ = .068, P = .123) (**Suppl. Fig. S1B**).

*Intravital Near Infrared (NIR) Imaging of MMP Activity*

One day prior to imaging, hair was removed from the lower extremities using hair clippers and a chemical hair removing agent. On the day of experimentation, mice were anesthetized, and a pre-injection image was acquired (Pearl Impulse, LI-COR, Lincoln, Nebraska). Mice were positioned supine with flexed knees and administered 4-μL injections of MMPSense into both joints (trans-patellar tendon) using a 33-gauge needle. After 2 hrs of normal cage activity, mice were re-anesthetized and re-imaged. Image analysis was conducted in a blinded fashion using ImageJ. Consistently sized circular regions of interest were placed over the knee joint, and the signal raw integrated density (RID) was quantified. RID of each limb was averaged across mice and analyzed in aggregate form or normalized to the contralateral limb.

*Micro Computed Tomography (μCT) Imaging and Analysis*

Femoral metaphyseal trabecular bone, femoral epiphyseal trabecular bone, and both medial and lateral femoral subchondral bone were analyzed using boneJ^31^, via the Matlab-ImageJ interface Miji^32^. Trabecular volumes were analyzed by bone volume fraction (BV/TV), bone tissue mineral density (TMD), trabecular thickness (Tb.Th), trabecular number (Tb.N), and trabecular spacing (Tb.Sp). Subchondral bone was analyzed by subchondral bone thickness (SCB.Th) and TMD. Derived morphometric parameters of both limbs in the ACLR group were then normalized to the mean value of Sham mice within each sex to calculate a percent change relative to the Sham group, which was compared between sexes and time points. To assess differences between Sham and ACLR limbs and between Sham and Contralateral limbs within each sex, aggregate non-normalized data were compared.

*Image Analysis of Na_V_1.8-Cre;tdTomato sections*

To quantify the proportion of Na_V_1.8+ surface area in synovium, ROIs of the medial synovium were manually outlined in each section, and Na_V_1.8+ signal was thresholded from background. The area of positive signal within each ROI was normalized to area, and the proportion of positive signal averaged in two sections spaced 80 μm reported for each limb. Further, the total number of Na_V_1.8+ SCB channels were counted in the medial femur and tibia. The average number of positive channels counted in two sections (80 μm apart) are reported. All quantifications were performed by a blinded observer (AMO).

*RNA Sequencing of Synovium and Bioinformatic Analyses*

Samples were homogenized in TRIzol (Thermo Fisher) using an automated tissue homogenizer (Precellys Evolution, Bertin Instruments). RNA was isolated using Qiagen RNeasy Micro kits (Qiagen), and all samples were confirmed to have RNA integrity number (RIN) values ≥ 6.5. Sequencing was performed using the DNA nanoball (DNB)-Seq platform (MGI Tech, BGI Americas Corp, Cambridge, MA), with 100 bp paired-end reads, >40M clean reads per sample. Reads were aligned to the *Mus musculus* reference genome (GCF_000001635.26_GRCm38.p6) using *Hisat^38^*. Aligned reads were mapped to reference genes using *Bowtie^39^,* and *RSEM^40^* was used to generate final read counts. Principal component analysis (PCA) of read counts (**Suppl Fig. S5C-D**) was used to identify outliers, and four out of 60 total samples were excluded as outliers. *DESeq2^41^* was used to identify differentially-expressed genes (DEGs), using the Wald test corrected with the Benjamin-Hochberg method. DEGs for all comparisons across all groups, sexes, and timepoints are listed in Supplemental Table 1. Pathway analysis was performed by submitting DEGs (P_adj_ < 0.05) for statistical enrichment or statistical overrepresentation testing using the Gene Ontology (GO) “biological process complete” annotation via PantherDB^42^.
