## Supplementary figures and images for "Sexual dimorphism of the synovial transcriptome underpins greater PTOA disease severity in male mice following joint injury"

### Fig S1

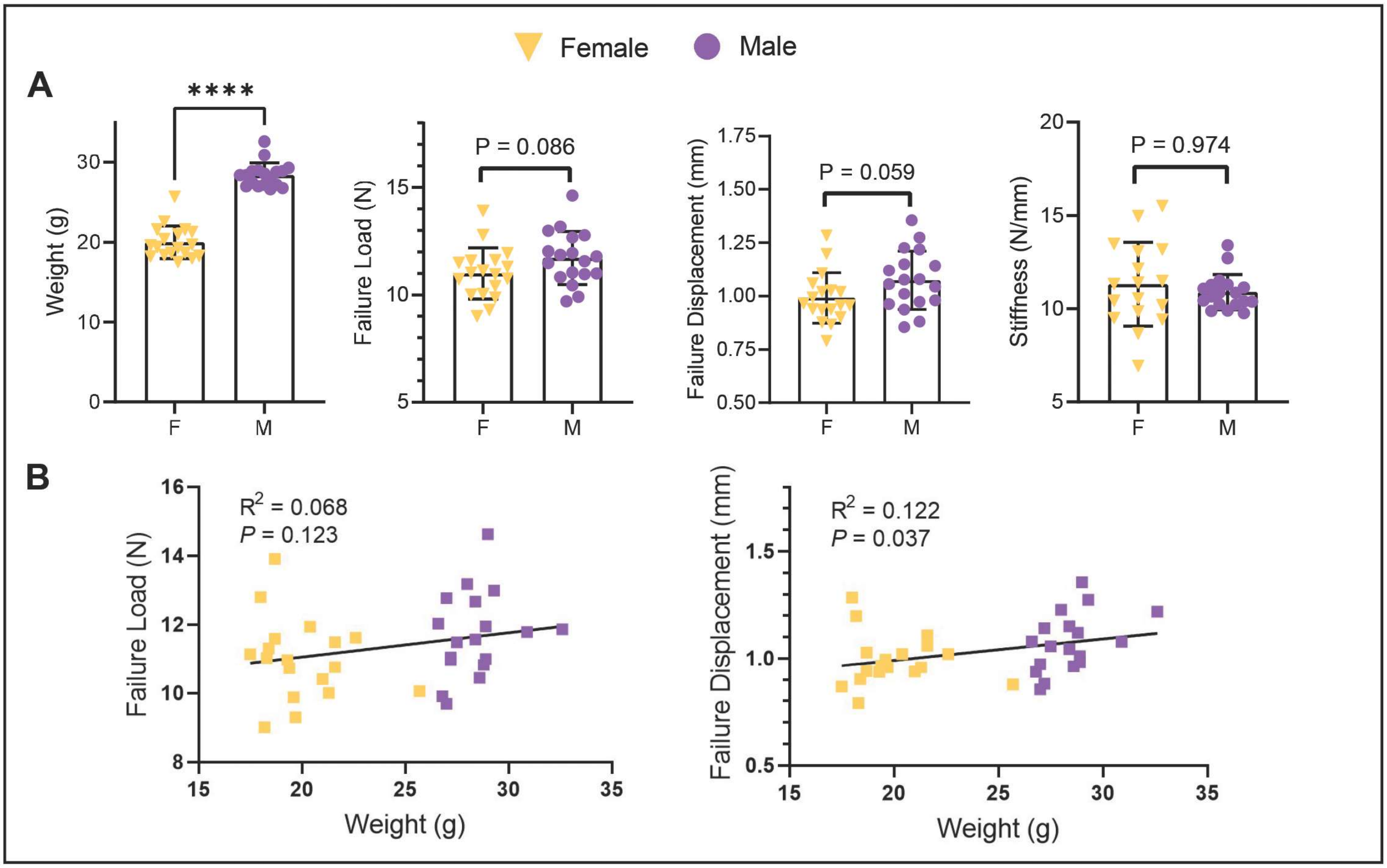

### Fig S2

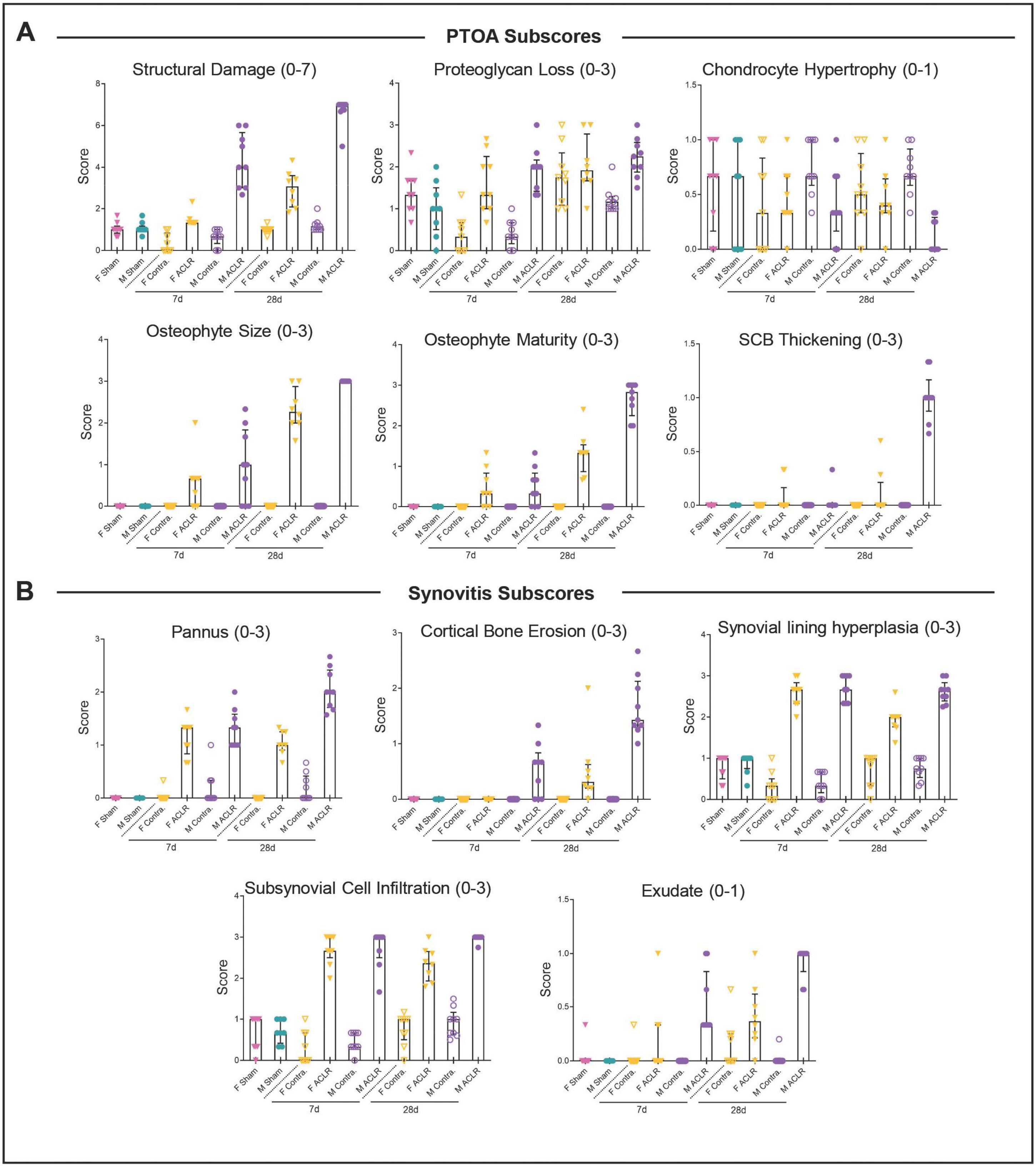

### Fig S3

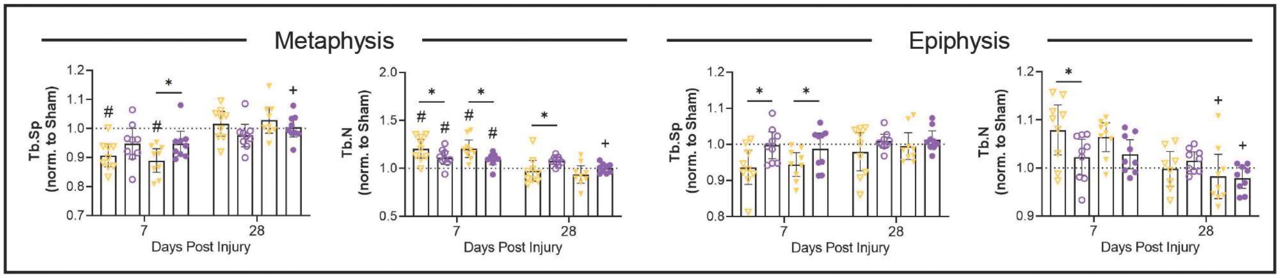

### Fig S4

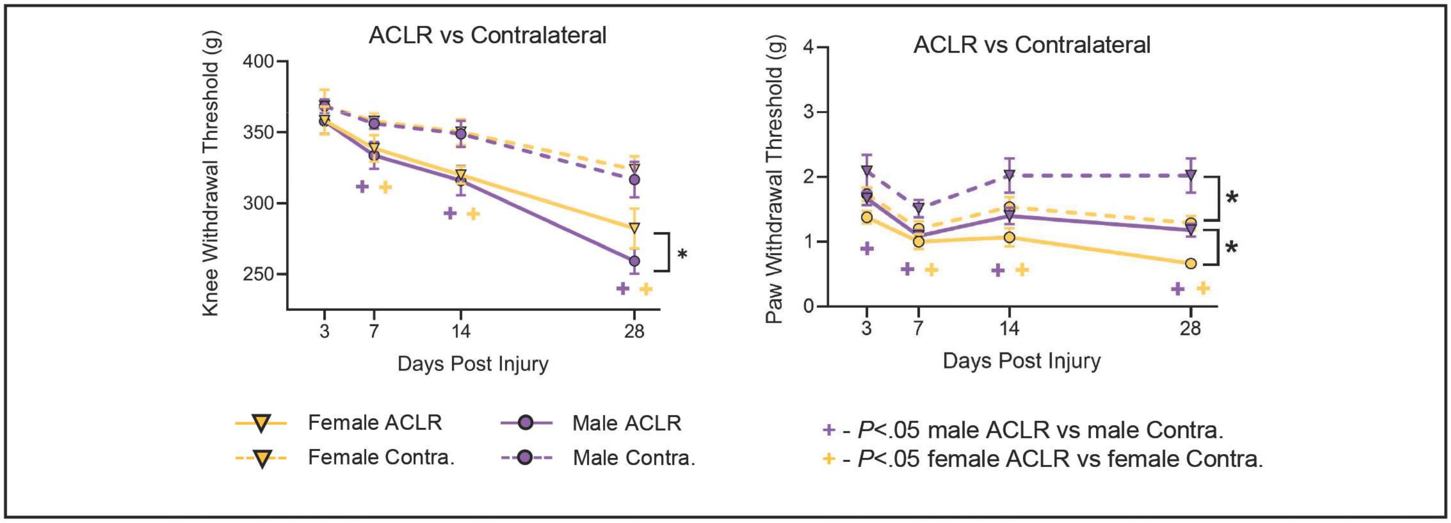

### Fig S5

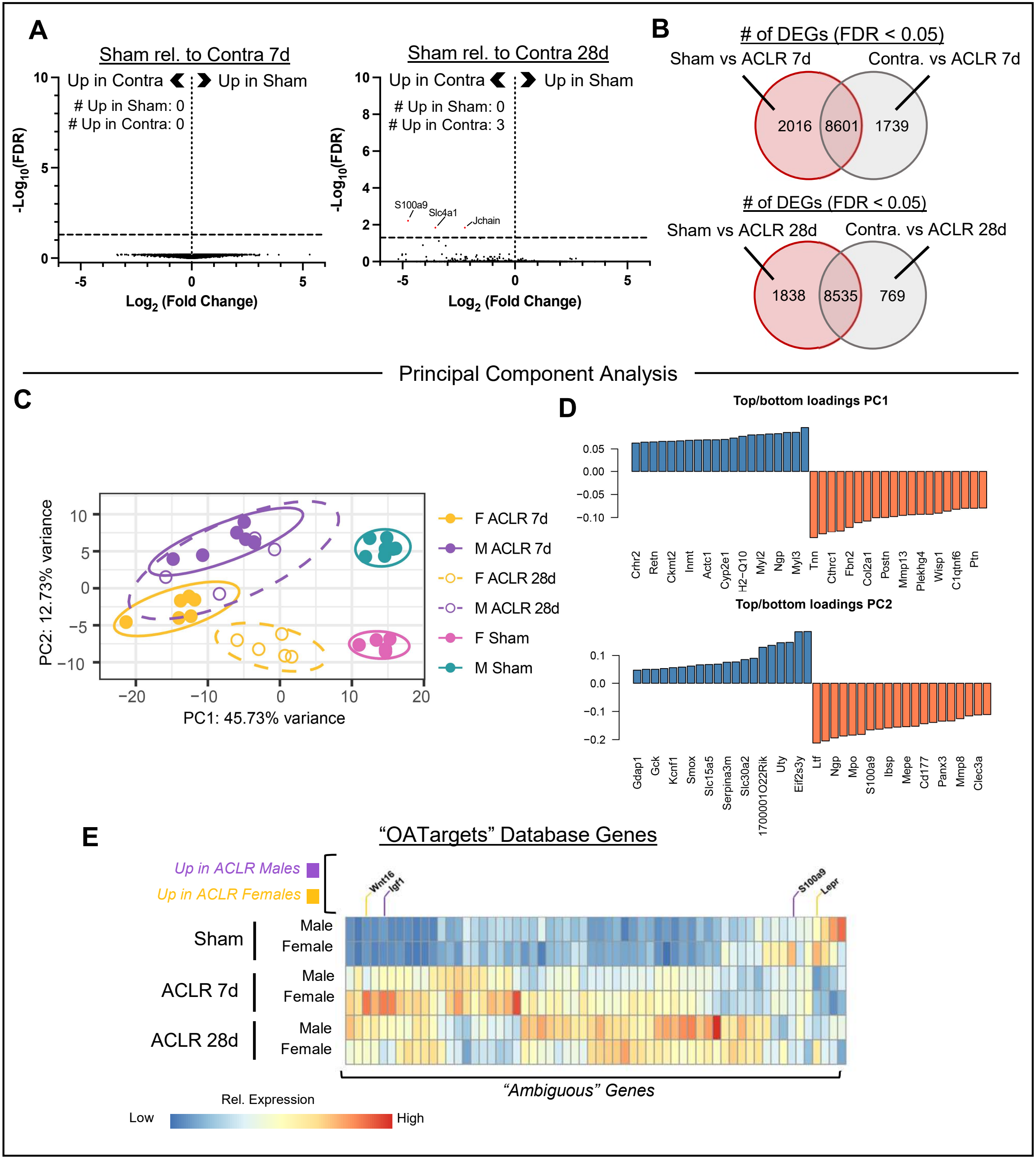
